# White-faced capuchins (*Cebus capucinus imitator*) exhibit selectivity for stone tools

**DOI:** 10.64898/2025.12.19.695614

**Authors:** Meredith K.W. Carlson, Brendan J. Barrett, Evelyn del Rosario-Vargas, Tamara Dogandzic, Margaret C. Crofoot, Nicolas Zwyns

## Abstract

Over the last few decades, the developing field of primate archaeology has used archaeological methods to reconstruct primate behaviors, especially tool use. We expand the possibilities of primate archaeology with an inverse approach: using living primates to address archaeological questions. In particular, we investigate tool selectivity, a phenomenon that not only represents a significant milestone for tool users, also for the emergence of stone tool technology in human evolution. Observing extant primate tool use can establish clear expectations for a percussion-based archaeological record across taxa. Here, we describe stone tools assemblages produced by two independent populations of white-faced capuchin monkeys (*Cebus capucinus imitator*) in Coiba National Park, Panama, and provide evidence of stone tool selectivity in both populations. We demonstrate that capuchins select for tool dimensions, and select different sizes for different tool functions. This selectivity analysis reveals that primate stone tool use results in assemblages of artifacts that are distinct from natural accumulations of unused material in the surrounding landscapes. These variations further suggest that stone tool use has a recognizable signature at the assemblage level, even where no lithic flaking occurs. The existence of such assemblage-level signatures offer new avenues for the archaeological identification of tool use both in unhabituated living primates and extinct taxa. The presence of selectivity in multiple capuchin populations suggests that, along with percussion, tool selectivity is ancestral to stone tool flaking, and is shared by phylogenetically distant branches of the primate tree.

## Introduction

The use of percussion tools has great phylogenetic breadth and temporal depth within the primate order. Percussion techniques are employed by all primate stone tool users, including *Pan* (Boesch & Boesch, 1983; Proffitt et al., 2022; Reeves et al., 2024), *Macaca* (4), *Sapajus* (5, 6), and *Cebus* (7–9). Many instances of primate percussion result in incidental flake production (10–12). These observations, along with arguments founded on cognition and social learning, support the assertion that percussion represents a precursor to the kind of stone knapping which characterizes the hominin archaeological record (13–15).

Among early human tool users, the targeted selection of materials is an indicator of increased investment in and knowledge of tool production (16, 17). In particular, selectivity for specific raw material types is a recognizable feature of Oldowan assemblages (18, 19). Selectivity among living primate tool users can clarify aspects of the capacity for and investment in tool use. However, there are also implications for archaeology itself: establishing the criteria through which tools are selected can set expectations for the composition of assemblages.

In the context of flake production, material selectivity indicates that toolmakers understand certain dynamics of the knapping process including conchoidal fracture, a principle of stone fracture mechanics essential to gain control of flake production (20). This means that in Oldowan contexts selectivity should – and often does – order itself around properties like cryptocrystalline structure and high silicate content, often summed up in terms of “high quality” and “low quality” materials (21–23). Yet, the characteristics that maximize technological efficiency are different for percussion tools than for knapped lithics. Although stone flakes are unintentionally produced by many primate tool users, sharp flakes are not the goal of these behaviors. Rather, in percussion the goal is to use the selected material to break a secondary object, rather than to control the fracture of material. As a result of these contrasting affordances, tool selectivity in percussion-focused assemblages will have a signature distinct from flake-based assemblages. Accordingly, raw material selectivity to maximize conchoidal fracture has not been observed in living nonhuman primate tool users (10).

The Lomekwian provides one model for how raw material selectivity might manifest in contexts that have a substantial percussion component and are not predominately composed of freehand flake production. In particular, the LOM3 assemblage demonstrates selection for cobble volume, not merely material quality (24). In particular, the largest locally available cobbles were selected for core reduction (25, 26). While subsequent Oldowan assemblages demonstrate more complex material selectivity with respect to fracture mechanics, cobble size distributions at LOM3 suggest that selection for more basic attributes, such as stone volume and weight, may be precursors to sophisticated selectivity criteria. These examples show that certain kinds of selectivity occur alongside the earliest stone knapping, but an increase in selectivity linked with the control of flaking events may emerge later. What remains unclear is whether raw material selectivity was induced by stone tool production, or whether it is a preexisting skill that was recruited from unmodified stone tool use.

In chimpanzees and *Sapajus* capuchins, selectivity has been observed for a number of tool characteristics, suggesting an ancestral condition for selectivity skills (1, 27). Among Taï Forest chimpanzees, tool users do not show preference for particular materials, but instead select hammerstones on the basis of proximity to the target, with material types reflecting local abundance (3). Original observations of tool selectivity in Taï chimpanzees included a preference for stone hammers in over wooden ones, and for heavier hammers when engaging in specific tasks such as pounding hard *Panda* nuts (1). In bearded capuchins (*Sapajus libidinosus*), evidence for tool selection according to weight has been mixed. Some studies have identified preferences among bearded capuchins for heavier hammerstones for processing hard palm nuts and lighter hammerstones for processing resistant resources (28, 29) . This result failed to be replicated in an experimental setting, in which most individuals were found to prefer nearby stones, regardless of stone weight (30). In a recent cross-population study, the tool weight selections of *Sapajus* tool users were explained by resource availability, rather than by properties of the target food (31). The existing literature suggests that while tool weight is likely to be an important selection criterion for primates using percussion tools, material selectivity remains to be established as a universal component of these technological behaviors.

If selectivity is present, we expect tool function to be the main driver of material selections for capuchins in Coiba NP, as percussion tool use in these populations involves processing of a broad range of resources with contrasting physical properties. Hammerstone percussion involves tasks ranging from breaking small, brittle snail shells to crushing large, fibrous almond and palm nuts, each of which could be optimized using stones of different weights. Capuchin stone manipulation mainly employs a power grip with either one or both hands, rather than the pad-to-pad precision grip characteristic of human tool use (32, 33). However, even among power-focused tasks there is variation in the amount of force required. The application of too much force can be detrimental to processing certain foods, for example by introducing non-edible materials such as shells, husks, or minerals that can damage tooth enamel (34) or causing processed foods to fly off an anvil and be stolen by a scrounging conspecific (35). Using too little force can result in failure, reduce energetic return rate, or increase exposure to the risks associated with tool use by increasing the processing time. These task-specific requirements and costs are expected to contribute to tool preferences where selectivity is expressed.

In this study we investigate whether white-faced capuchins (*Cebus capucinus imitator*) in Coiba NP, Panama exhibit raw material selectivity during stone tool use, and whether tool choices are functionally specific. In the absence of active selection, the properties of used tools should reflect those of stones that are available near food-processing sites. In contrast, general and task-specific selectivity for particular stone properties will produce identifiable assemblages of unmodified stones which are distinct from local material abundances.

## Results

### Assemblage description

We identified a total of 528 hammerstones during site surveys, 187 on Coiba and 341 on Jicarón. Anvils at tool sites were a mixture of stone, driftwood, and trunk bases, buttresses, and branches of live trees. The majority of hammers were associated with stone anvils (Jicarón: 65.7%, n = 224; Coiba: 92.5%, n = 173). Most identifiable hammers were recovered intact (88.4%, n = 467), however surveys also identified fragmentary hammers, including complete refits (5.9%, n = 31) and partial refits (5.7%, n = 30). Hammerstones were of various local rock types, including basalt, limestone, fine-grained sandstone and occasionally quartzite. On both islands, stones are primarily rounded cobbles from secondary sources in streambeds and on beaches.

Due to our protocol for identifying tool use, all hammers were, by definition, associated with an anvil. However, some sites contained multiple hammers. Similarly, while all tool sites had evidence of at least one type of food debris, some sites had evidence of more than one resource. The most common debris associated with tool sites on Jicarón was *Terminalia catappa* exocarps (76.5%, n = 261). Other associated debris included *Coenobita compressus* (49.9% of hammers, n = 170), coconuts (26.1% of hammers, n = 89), and *Gecarcinus quadratus* crabs (7.3% of hammers, n = 25). On Coiba, the most common debris associated with tools were shells of freshwater Nerite snails (*Clypeolum latissimum)* (47.6%, n = 89) and *Astrocaryum standleyanum* nuts (35.3%, n = 49). *Cocos nucifera* and *Bactris major* fruits were also found at low frequencies (< 3%) at tool sites on Coiba.

### Hammerstone selectivity

We compared the assemblages of utilized tools by function with locally available, unused raw materials (Table S2). The heaviest hammerstones are those used for processing *Terminalia* nuts (median weight 675 g) and *Astrocaryum* palm nuts (median weight 870 g). Hammers used to process *Terminalia* included the single heaviest hammer recorded, at 3000 g. While nut-processing tools were on average heavier than raw materials, these tools were not tightly constrained in weight, ranging from 56g to 3000g. Hammers used for processing small invertebrates were less distinct in weight from raw materials. *Coenobita compressus* crab hammers (Jicarón) had a median weight of 400 g and Nerite snail hammers (Coiba) had a median weight of 200 g.

Posterior distribution analysis shows differences between cobble weight by both function and tool/raw material status. For the Jicarón assemblage, our model predicts hammerstones used to process both *Terminalia* nuts and hermit crabs to be heavier than raw materials, with *Terminalia* hammerstones predicted to be larger overall (Table S3). Similarly, for the Coiba assemblage, tools used to process *Astrocaryum* nuts are predicted to be heavier than local raw materials. Tools used for processing *Clypeolum* snails are predicted to be lighter than raw materials (Table S4).

When comparing the contrasts of posterior estimates of mean metric dimensions of tool categories relative to raw material, we found no to very little overlap of posteriors with zero (Figure 4). This indicates that there is a signature of selectivity for length, width, thickness, and weight. Weight showed the largest effect on tool status on both Jicarón and Coiba. On Jicarón, *Coenobita* crab hammers and *Terminalia* nut hammers were larger across all dimensions than were raw materials. On Coiba, *Clypeolum* snail hammers were smaller across all dimensions, and *Astrocaryum* hammers were larger across all dimensions.

To assess selectivity related to specific functions, we modeled weight and linear dimensions among tools used for different functions. We limited our analysis to the most common resources found at tool sites, excluding resources that were found in less than 10% of sites. On Jicarón, the resources included in the analysis were *Terminalia* nuts and *Coenobita compressus* hermit crabs. On Coiba, *Astrocaryum standleyanum* palm nuts and *Clypeolum* snails were included. Coconut was excluded from the analysis, as camera trap video from major tool sites suggests that coconuts are most often processed directly on the anvil, without the use of a hammerstone – hammerstones appear to be used on shards of coconut shells to help access fruit flesh during secondary processing.

To compare selectivity along metric dimensions for hammerstones relative to raw materials, we modelled the population distributions of hammerstones (by tool function) and raw materials (Figures 2, 3). Where prediction estimates overlap for a particular dimension (i.e. weight), tools are selected for use proportional to the abundance of same-sized raw materials in the local environment. In other words, there is no selectivity beyond natural abundance. Where posterior predictions are lower or higher for tools relative to raw materials, this indicates that tools are being selected at a rate either lower or higher than their expected distribution in the population. These are interpreted as signatures of selectivity.

**Figure 1.**
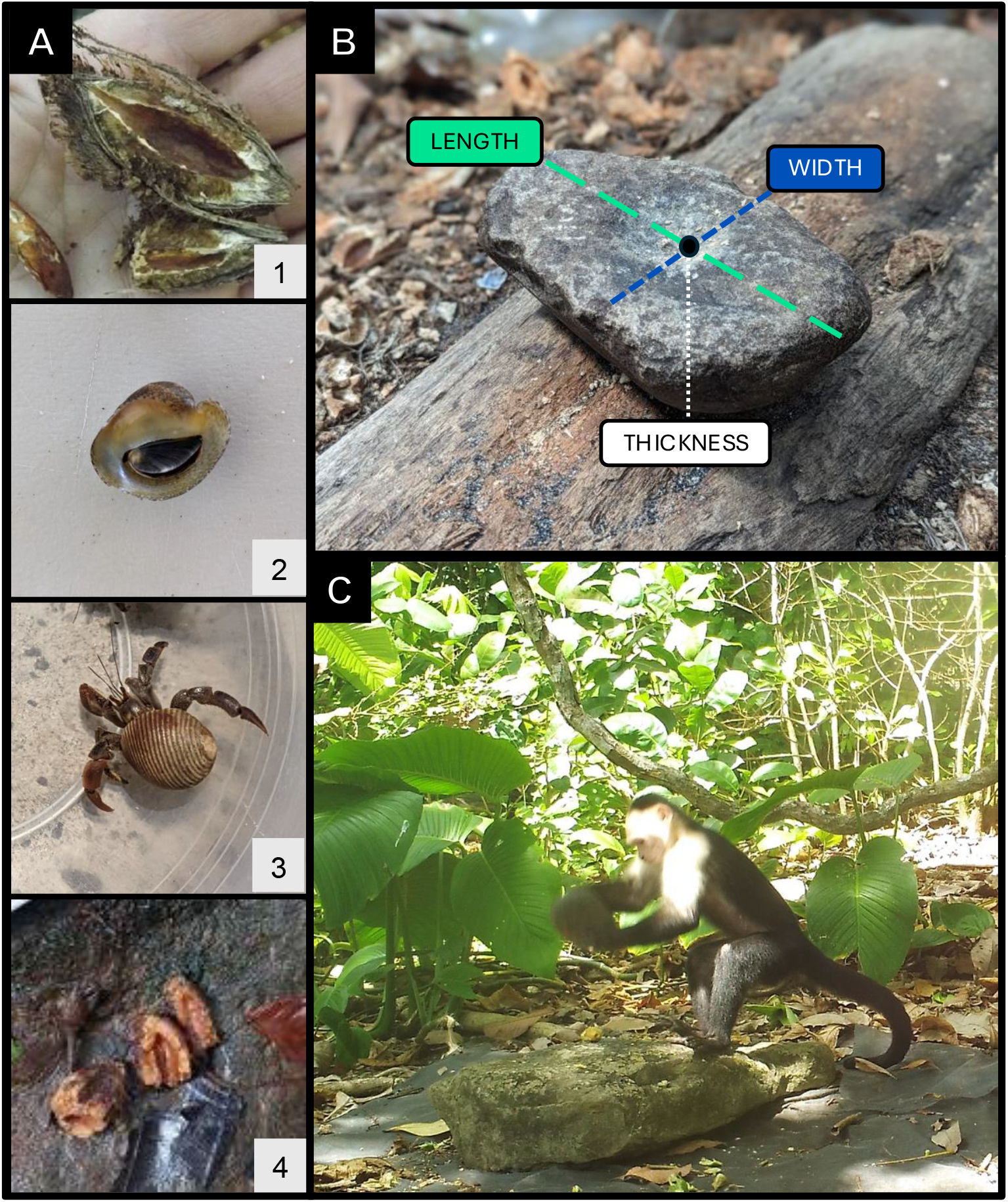
A) Resources processed by capuchin tool users: (1) *Terminalia catappa*, (2) *Clypeolum latissimum*, (3) *Coenobita compressus*, (4) *Astrocaryum standleyanum*. B) Linear measurements of hammerstones. C) White-faced capuchin percussive tool use on Jicarón via camera trap.

**Figure 2.**
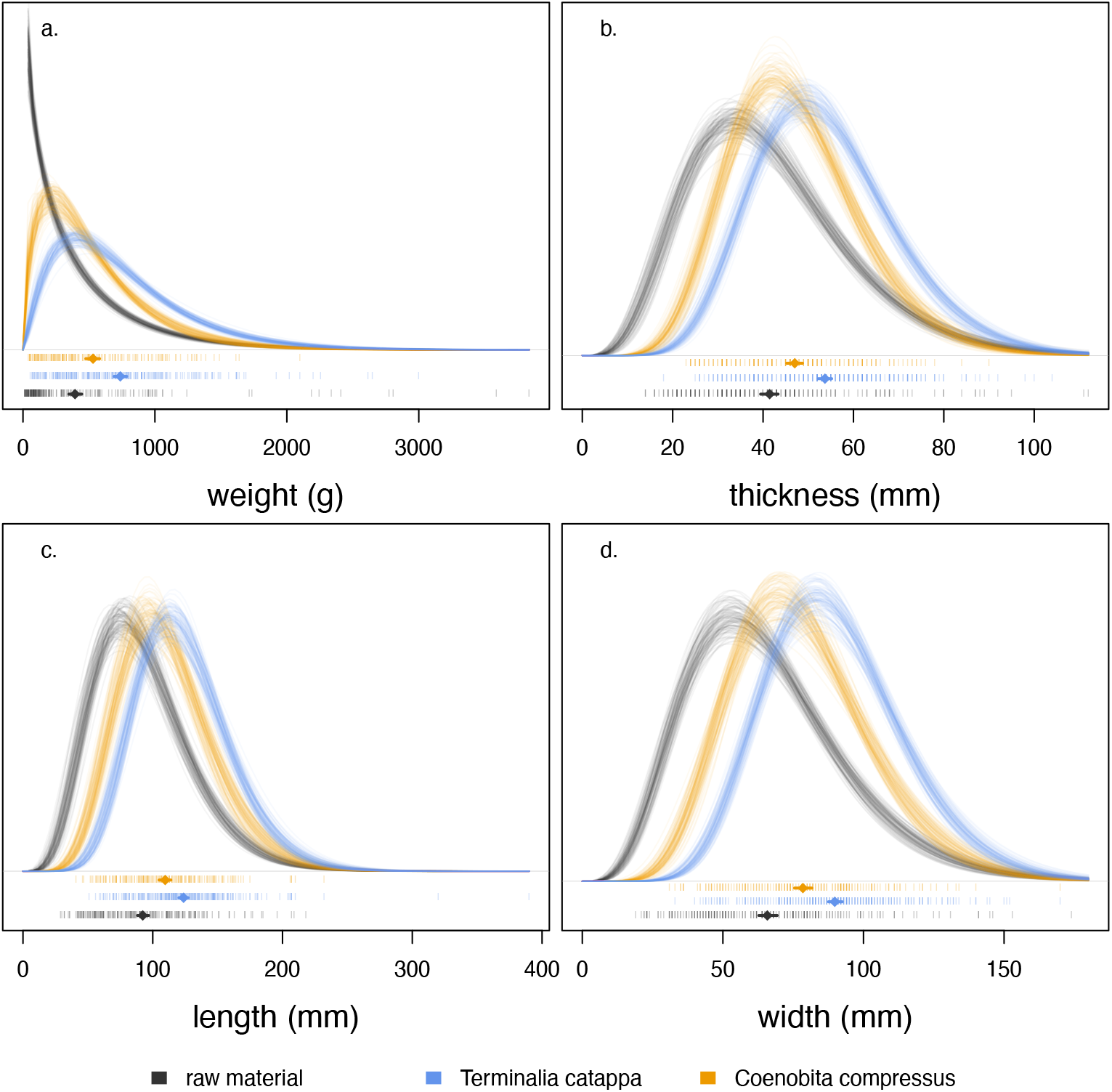
Posterior distributions comparing weight and linear dimensions of raw material (grey) to utilized tools for Jicarón materials. *Terminalia* hammerstones (blue), and *Coenobita* hammerstones (orange). Lines show 100 marginalized posterior predictions drawn from a gamma distribution for each dimension/category combination. At the bottom of each panel are diamonds indicating the posterior mean for each dimension estimated from each model, and a horizontal line corresponding to the 89% HPDI, overlaid over raw data (vertical dashes).

**Figure 3.**
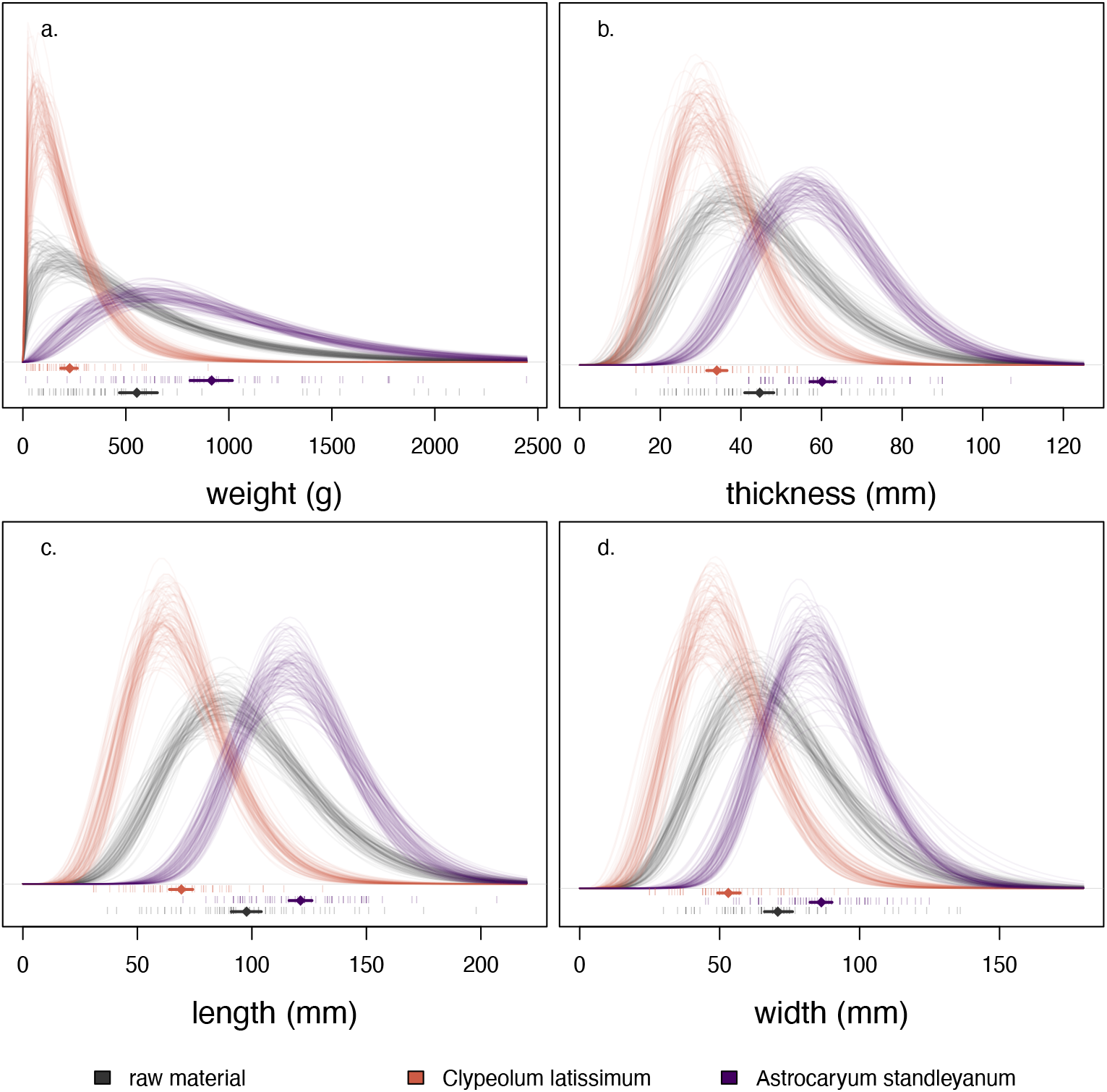
Posterior distributions comparing weight and linear dimensions of raw material (grey) to utilized tools for Coiba materials. *Astrocaryum* hammerstones (red), and *Clypeolum* hammerstones (purple). Lines show 100 marginalized posterior predictions drawn from a gamma distribution for each dimension/category combination. At the bottom of each panel are diamonds indicating the posterior mean for each dimension estimated from each model, and a horizontal line corresponding to the 89% HPDI, overlaid over raw data (vertical dashes).

**Figure 4.**
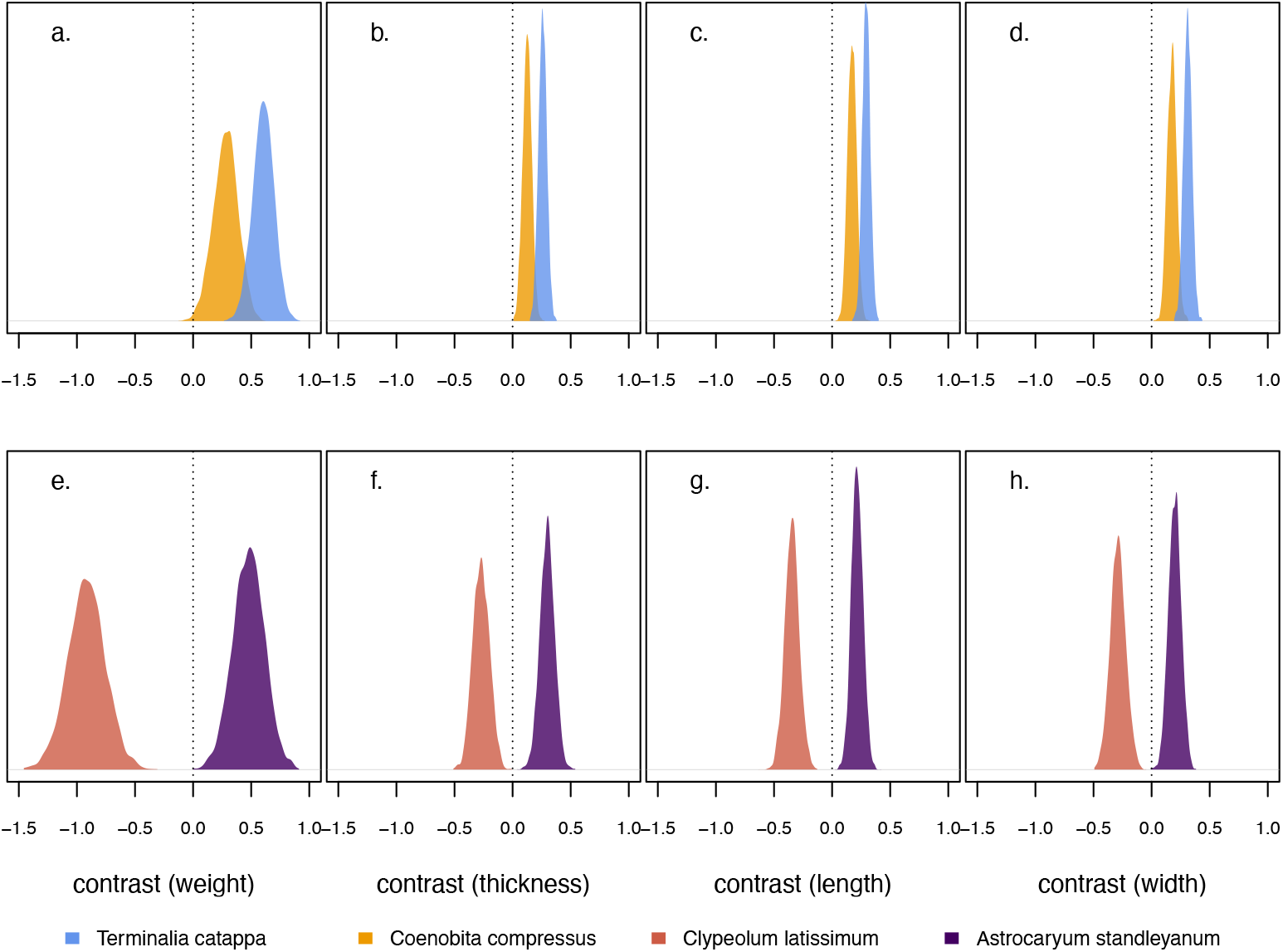
Estimated posterior contrasts of each mean tool dimension (weight, thickness, length, width) for individual tool functions, relative to raw materials. Negative contrasts indicate a smaller size for the tools than raw materials, whereas positive values indicate a larger size relative to raw materials. Dashed line lies at zero, indicating equivalence. (a-d) Tools from Jicarón, (e-h) Tools from Coiba.

On Jicarón, there is selectivity against *Coenobita* hermit crab hammers less than ∼200g in weight, and selectivity for hammerstones between ∼200 and 1600g, relative to the availability of such materials. For hammerstones used on *Terminalia* nuts, there is selectivity against hammers less than ∼400g, and selectivity for hammers between 400 and 1800g. On Coiba, selection of *Clypeolum* snail hammers is biased toward stones < ∼350g and biased away from stones > 450g. For hammerstones used on *Astrocaryum*, there is selectivity for hammers between ∼500 and 2200g.

## Discussion

With recent increased interest in the field of primate archaeology, researchers have called for more comprehensive investigation of primate behavior through archaeological methods (39). Here, we respond both to this call and also its inverse: using primate data to address archaeological questions, broadly construed. Primate archaeology has the potential not only to address behavioral questions, but also to generate a framework for interpreting the archaeological record. A shared obstacle to both primate archaeology and Plio-Pleistocene archaeology is the (potential) simplicity of the toolkits used. While many primate tool users produce incidental flaked lithics (2, 10, 11, 40–42), the primate archaeological record is not composed of the kind of intentional, and therefore highly conspicuous, artifact reduction that defines the majority of identifiable hominin assemblages. Further, applying use-wear analysis in order to identify contemporary primate tools may only be effective in cases where the tool use is particular intensive, and not all tools will be identifiable in this way (12, 43). In addition to planning and investment, understanding tool selection establishes expectations for the composition of low-visibility artifacts and assemblages.

### *Task-specific selectivity in* Cebus

Overall, our findings demonstrate that task-specific selectivity in tool use exists in the genus *Cebus*. White-faced capuchins are able to tailor their choice of stones to meet the requirements of different pounding tasks. While there is overlap in the characteristics of individual hammerstones and individual raw material cobbles, used tools and raw materials are distinguishable on the assemblage level. Raw material choices are restricted by function-specific selectivity for different food-processing tasks. Most notably, heavy tools were used for nut-cracking (*Terminalia catappa* and *Astrocaryum standleyanum*), and light tools were chosen for processing *Coenobita compressus* and *Nerita* and *Clypeolum* snails. These selection criteria make for tools that are particular suited to the task at hand. *Astrocaryum* and *Terminalia* nuts are hard-shelled, durable, and somewhat elastic, requiring the application of high force to break open the endocarps. *Terminalia* nuts are also highly fibrous, so episodes of hammerstone processing are usually completed by capuchins tearing open the exocarp with the hands. Unlike with processing of nuts, the application of too much force in opening invertebrate shells can result in the destruction of the food within, rendering it largely inedible. The presence of a clear upper limit on tool size and weight, relative to body size, for *Clypeolum* snail and *Coenobita* crab tools, but not nutcracking tools, indicates that capuchins are not only selective about their tools, but are able to make selections that maximize food return while minimizing energetic expenditure. When required for nutcracking, capuchin use some of the largest and heaviest cobbles available, but avoid doing so for other tasks. This maximization of tool weight for high-force tasks, within the constraints of our sampling criteria, indicates that selectivity is related to function and efficiency, rather than the constraints of capuchin physical abilities. The use of heavy hammerstones increases the force applied to crack or crush tough food items. However, manipulating these heavy hammerstones comes with an energetic cost. Hammers used to process *Terminalia* sometimes exceeded 3kg. This represents a substantial investment for capuchins, as adult male average body weight is 3.7-3.9kg, and capuchins from Coiba are reported to be slightly smaller than their mainland counterparts (44). Given that large hammers approach and may exceed 75% of adult body weight, it is particularly notable that capuchins select these stones for nut-cracking tasks – tasks to which heavy hammers are especially suited.

An outstanding question is whether capuchins exhibit selectivity solely for cobble weight, or whether they account for other dimensions when choosing hammerstones from among available raw materials, and if this varies across foraging contexts. Palm height and grip may place limits on the dimension of thickness, requiring particular dimensions for great control during percussion. A similar constraint may exist for width, in which a hammer that is too wide (or narrow) is more difficult to control or target. Establishing selectivity for individual dimensions is a challenge, as linear dimensions scale allometrically with mass. We envision two potential routes to address this question. The first is to perform behavioral experiments with wild capuchins, in which material weight is controlled and linear dimensions are experimentally manipulated. Such experiments would also have the benefit of isolating some of the behavioral cues that capuchins may use in their material selection decisions. The second route for identifying selectivity for particular dimensions would use covariance analysis. In raw materials, we would predict strong correlations between weight and all linear dimensions. However, selectivity for a particular dimension (i.e. weight) or dimensions (i.e. weight and thickness) would be expected to lead to a lower correlation between the selected linear dimension and the non-selected dimensions, when compared to raw materials. Analytical solutions, similar to those used in population genetics to identify natural selection, might be applied to this context in order to determine whether selectivity occurs with regard to one or multiple metric dimensions.

We describe a kind of hammerstone selectivity that does not require long-distance transport of stones – white-faced capuchins do not appear to transport tools further than a few meters, and may actively avoid transport. Resources which capuchins use tools to open are common in rocky intertidal, coastal, and riparian zones – areas where lithic material is abundant, but where the persistence of individual stones may be limited by tides and floods. Despite this, utilized tools show a signature of selectivity. Tools represent a biased subset of the variation present among stones from nearby streambeds and beaches. Further, tool sites are highly concentrated within the study area: sites are localized to small (approximately 2 km) stretches of coast and river on each island. This may indicate that, within the home range of groups that have a tradition of tool use, the locations of tool use sites are linked to areas where suitable stone material is readily available. Sites may also concentrate around the locations of food resources, although the precise factors driving the distribution of tool sites within the home ranges of tool-using groups still need to be determined.

### *Defining the* Cebus *toolkit*

Through the assemblage analysis, we have identified the composition of the *Cebus* stone toolkit. Our results suggest that, despite its relative simplicity, the toolkit in each of these two populations contains more than one tool type. While *Cebus* stone tools are not intentionally modified, the selection of different properties for different tasks leads to the accretion of assemblages composed of groups of tools with distinct qualities and, arguably, distinct functions. Small invertebrate hammers function mainly as “fracturing” tools, while large nut hammers function as “crushing” tools. Furthermore, capuchins manipulate crushing hammers and fracturing hammers in distinct ways. Video footage from these sites has shown that capuchins primarily use a two-handed grip when pounding nuts with stone tools (Video S1). However, Nerite snail and *Coenobita* crab hammers are often used with a single-hand grip, in keeping with the limitations of the small average size of both tool and target in these cases (Video S2).

### Paleoanthropological implications

In the archaeological record, selectivity is a milestone of investment and skill in hominin tool production (17, 19). Targeted material choices co-occur with the early knapping, where the preferential selection of cryptocrystalline, silicate materials offers greater control over stone flaking. Beyond selection for typical “high quality” rock types, the early archaeological record for hominins, as well as data from extant primate tool users, suggests that weight may be important for percussion activities. We establish that selectivity for stone weight is present in capuchin percussion tool use, suggesting that the skill for appropriate material choice of this kind may precede the development of intentional modifications of stone through knapping.

These results show that stone tool selectivity is likely to be ancestral to stone knapping for hominins, both by dint of being phylogenetically widespread among primates and having a low barrier to independent, convergent innovation. We provide new evidence that the capacity for tool selectivity can be found broadly in platyrrhines, as well as hominoids. The presence of similar, task-specific selectivity in two independent capuchin populations isolated for thousands of years offers support for the idea that certain fundamentals of stone tool use may represent “latent solutions” which can be reinvented without the need to invoke high-fidelity social learning (45). While selective pressure for efficient material choices undoubtedly increased as the complexity and dependence on lithic technology increased throughout human evolution, targeted and effective selectivity for tools is likely to be a pre-existing capacity for tool using primates.

### Implications for identifying unmodified assemblages

The observation of capuchin selectivity is also significant because it allows us to describe the nature of assemblages produced by percussion. Capuchin tool choices create assemblages that are distinct in weight and size from nearby sources of stone. Such distinctive assemblages are identifiable markers of tool use, even in the absence of tool manufacture and organic debris from tool-aided foraging. This archaeological visibility has implications not only for primate archaeology, but for the identification of early hominin tool use. The earliest lithic assemblages currently described contain a higher ratio of hammerstones to cores and flakes than later, more developed Oldowan assemblages (25, 46). Heavy-duty percussion continues to play an important role in some lithic assemblages through the Oldowan and Acheulean (47–50). Further, unmodified, transported stones (“manuports” and “utilized material”) have been identified as artifacts in paleolithic assemblages for decades (51). However, the interpretation of these stones as true artifacts has been challenged (52). At present, manuports and utilized pieces have only been identified in assemblage that contain significant flaked components, due to the difficulty of establishing use in the absence of modification. It is therefore significant that in the capuchin context presented here assemblages of stone tools which have been used, but neither intentionally modified nor accidentally flaked, have distinct signatures. Recent work has demonstrated the overwhelming likelihood that nut-cracking activities can be identified through the presence of flaking, if suitable raw materials are used (10). Our findings, which establish an assemblage-level signature of tool use in the absence of flaking, suggest that tool use in both extant primate and hominin contexts might be recognized in a broader set of circumstances than has previously been explored. Where identification via use wear or either intentional or incidental modification is not possible, our results indicate that comparison of materials with the surrounding landscape may allow tool use to be identified. As the earliest dates of the hominin archaeological record are pushed deeper into the past, it becomes increasingly clear that early assemblages contain significant percussion components, including unmodified elements. The identification of selectivity among capuchin tool-users broadens our ability to trace technological evolution and to interpret tool use in the deep past.

## Materials and Methods

### Study area

Our field study was conducted at two locations within Coiba NP where populations of white-faced capuchins (*Cebus capucinus imitator*) use stone tools. Coiba NP is located off the Pacific coast of Panama, at a distance of approximately 22 km from the mainland. The park encompasses a continental archipelago on the Cocos plate, which was isolated from the mainland 15,000-18,000 years ago, after the last glacial maximum (53). The geology of the archipelago is primarily volcanic: the islands are largely of basaltic lava composition, and are also made up of breccia and limestone (54, 55). The first capuchin tool use location is along the western coast of the island of Jicarón, where tool use was first described for this genus (8). The second location is in the northwest of the island of Coiba (9). The islands of Coiba and Jicarón are separated by 6 km at the closest point, and the two study sites are separated by a distance of approximately 38 km. The tool using populations on each island are isolated from one another, and it is believed that tool use is an independent innovation within each (9). The two islands therefore present the opportunity for a natural replication experiment in tool choice and selectivity.

The majority of forest in Coiba NP is primary growth (56). Coiba floral and faunal communities exhibit reduced alpha diversity compared to the mainland of Panama, but show high levels of endemism (57). While we estimate the population density of white-faced capuchins to be high on Jicarón, few capuchin groups on the island use tools. Stone tool use has been identified only among groups living near a ∼2 km coastal stretch on the western part of the island. Preliminary surveys across the entire coast of Jicarón and parts of its interior show no evidence that other capuchin groups beyond this area use tools. Tool sites with artifacts occur primarily in the intertidal zone and along permanent and seasonal streams in the coastal forest on Jicarón, where both raw materials and tool-processed food sources are abundant. Survey on the island of Coiba has been less extensive, and stone tool use has been identified at a locality along Rio Escondido in the northwest of the island.

The capuchin toolkit on both Coiba and Jicarón consists of hammer (active) and anvil (passive) elements used in tandem to process foods (Figure 1c). Hammers and anvils are frequently stone, but wood tools – primarily anvils – are also used. We limit our analysis here to stone hammers. Capuchins in Coiba NP use percussion tools to process *Terminalia catappa* nuts (English: sea almond; Spanish: almendro de playa), *Bactris major* fruit (English: Bactris palm; Spanish: caña brava), *Cocos nucifera* (English: coconut; Spanish: coco), *Coenobita compressus* (English: Ecuadorian hermit crab; Spanish: hermito), *Nerita scabricosta* (English: marine Nerite snail; Spanish: caracol nerita acanalado), *Clypeolum latissimum* (English: freshwater Nerite snail; Spanish: caracol nerita marino), and *Gecarcinus quadratus* (English: Halloween crab; Spanish: cangrejo de tierra rojo) (8, 9). During this study, we found evidence of additional stone tool functions not previously reported for this species – primarily the use of tools to process *Astrocaryum standleyanum* nuts (English: black palm, chumba-wumba; Spanish: chonta) (Figure 1a).

### Tool site identification

On Jicarón, capuchins use a series of clustered streambeds (permanent or seasonal) and the adjacent beaches for tool use. Capuchins frequently make use of the intertidal zone during foraging at this location (58). On Coiba, all identified instances of tool use occur along one permanent river and the neighboring beach near its mouth. We concentrated our surveys to identify tool sites around these temporary and permanent water courses, both because of capuchin foraging activity in these areas and the occurrence of appropriate stone for use as hammers and anvils. Streambeds and beaches provide opportunities for capuchins to encounter rounded stone cobbles that have eroded out of their original deposits. The extraction of stone materials directly from these primary sources has not been observed.

On Jicarón, opportunistic surveys to record tools focused on streams where tool use had been previously identified. These surveys consisted of walking from the coast upstream until impassable barriers (e.g. waterfalls or cliffs) were reached. We also conducted surveys of the beach in the intertidal zone along the coast where these streams terminate. On Coiba, surveys were conducted along the banks of Rio Escondido and within the course of the river. Surveys were also conducted in the intertidal at the termination of the river. Site surveys were conducted between July 2021 and December 2024.

For archaeological comparability, we use the term “site” to refer to the unit associated with an individual anvil. Because this definition centers on toolkits and not occupation zones or group ranges, sites typically encompass an area of no more than 4m^2^. Capuchin tool use sites are identified by the concurrence of three elements: an anvil, one or more hammers, and the debris of resources processed on the anvil. Food debris may be found on or adjacent to the anvil, or embedded on the hammerstone. The association of these three elements is sufficient to identify tool use, although macroscopic use wear (e.g. pitting, striations, discoloration) is also evident on some hammers and anvils, providing further evidence of use. A small number of hammers have deep pitting, similar to that described at other nonhuman primate sites. Some stone anvils also have shallow macroscopic pitting, but major surface modifications through deep anvil pitting have not been observed.

For the analysis, the tool function was defined by the resource type or types associated with the site. Sites vary greatly in the volume of food debris accumulation. While some sites are the product of a single episode of tool use, others presumably represent time-averaged activity, and are composed of debris from multiple independent tool-using events. As a result, some sites contain multiple types of debris. For sites with debris accumulation from multiple event and/or resources, all types of resources represented by the debris were included.

For each site, we recorded metrics of the hammer(s), the GPS location and accuracy of the site, and the type(s) of food debris associated with the site. Survey data were digitally recorded using a KoboCollect form on an Android smartphone.

### Hammerstone metrics

Capuchins do not intentionally modify the cobbles they use as hammerstones. As a result, individual artifacts can rarely be identified out of context. We include here only hammers that were found in context at an identifiable tool use site. We collected standard lithic measurements in the field, describing the length, width, thickness, and weight of hammerstones. We used manual calipers (1 mm accuracy) to measure length (maximum dimension), width (maximum perpendicular to length), and thickness (intersection of length and width) of each tool (Figure 1b). To collect weight, we used both a digital kitchen scale regularly checked for calibration (1 g accuracy) and a 2000 g Pesola Hanging Scale (Pesola AG, Chur Switzerland ; 5 g accuracy). In some cases, sites contained hammer fragments that could be refit. Where a complete refit was possible, the weight and dimensions of the original hammer were reconstructed and included in the analysis. Data on incomplete hammer fragments were collected but not included in the analysis.

### Raw material survey

Surveys of raw material on Coiba and Jicarón were conducted in order to establish a baseline of the characteristics of stones available to capuchin tool users. We conducted transect samples of stone cobbles in the intertidal zone and streambeds on both islands. In the intertidal, we sampled stones along two transects: a high-tide transect located at the vegetation line, and a mid-tide transect located between the high and low tide points on the day of sampling. The method used for sampling cobbles was adapted from the Wolman method (59). In this method, cobbles are selected blindly at regular intervals along a transect. In accordance with the small area of capuchin sites, we sampled raw materials at 1 m intervals. We limited our sample to stones of dimensions that could realistically be moved or manipulated by capuchins. We excluded stones with length (maximum dimension) smaller than 15 mm and larger than 400 mm. We sampled a total of 202 cobbles on Jicarón between July 2021 and July 2022. On Coiba, we surveyed raw materials during the July 2022 and December 2024 field seasons, collecting a total of 63 cobbles.

### Models of hammerstone selectivity

For tool sites on both islands (Coiba and Jicarón), we aimed to determine whether capuchins select hammers of particular weights and linear dimensions, with respect to particular tool functions. These functions were defined by the two main processed materials on each island: Jicarón: *Terminalia catappa* and *Coenobita compressus*; Coiba: *Astrocaryum standleyanum* and *Clypeolum latissimum*. Tool categories for each function were compared with raw materials from their respective island.

To analyze these data we used a series of Bayesian generalized linear mixed effect models fit using the rethinking package v. 2.40 (60), a wrapper to the MCMC sampler Stan (61) in R v.4.40 (62). As our outcomes are all measurements greater than 0, we chose a gamma distribution as our outcome with a log link. We estimated a unique mean (μ) and scale (φ) parameter for all four metric dimensions (weight, length, width, thickness) φ for all raw material and tool category (by tool function), yielding 24 unique mean and scale parameters. To compare differences in posterior predictions of the means of metric dimensions, we calculated contrasts of each tool category relative to its reference raw material category (Figure 3).

Estimating unique mean and scale parameters for each category permits us to model the entire population distribution of each raw material and tool category. This was necessary to assess potential selectivity, as selectivity can manifest as a bias away from or toward extreme values for dimensions. Similar to balancing or disruptive natural selection, such selectivity would not shift the mean, despite selection occurring. We accomplished the comparison of differences between posterior distribution of tools relative to raw materials by plotting posterior predictions of the plausible populations of tools and raw materials along all four metric dimensions, using both the shape and scale parameters.

For our models, we performed prior predictive simulations to ensure our priors specified a wide range of biologically plausible values, and ensured models converged by evaluation of R-hat values and trace plots. The models utilize 4 chains, which ran for 200 iterations each, half of which were used for warmup. We also performed an analysis that evaluated whether the timing of each data collection cycle affected variation in our estimates of metric dimension and found little variation across trips (Table S1). Accordingly, we proceed with the simpler Bayesian GLMs. Reproducible code and data for analyses are publicly available (https://doi.org/10.5281/zenodo.17643906).

This study was conducted with permission from the relevant authority, Ministerio de Ambiente, Panama, and was minimally invasive with regard to animal subjects. No animal care and use protocol was required for this study.

## Author Contributions

**MKWC**: Conceptualization, data curation, formal analysis, funding acquisition, investigation, methodology, project administration, visualization, writing – original draft, writing – review & editing

**BJB**: Conceptualization, data curation, formal analysis, funding acquisition, investigation, project administration, visualization, writing – original draft, writing – review & editing

**EDRV**: investigation

**TD**: Conceptualization, investigation, writing – review & editing

**MCC**: Funding acquisition, writing – review & editing

**NZ:** Conceptualization, writing – review & editing

## Acknowledgments

This research was supported by a Leakey Foundation Grant (F202310477) awarded to MKWC. It was also supported by a Packard Foundation Fellowship (2016-65130), a grant from the National Science Foundation (NSF BCS 1514174), and by the Alexander von Humboldt Professorship endowed by the Federal Ministry of Education and Research awarded to MCC. Permission for this study was obtained from the Ministerio de Ambiente, Panama (scientific permit no. SE/A-37-17, SC/A-23-17, SE/A-98-19, SE/A-6-2020, ARB-158-2022, and corresponding renewals and addenda). We sincerely thank Pedro Castillo, James Chaves, Bradley Christin, Mackenzie Godfrey, Kipp Godfrey, Zoë Goldsborough, Odd Jacobson, Korayma Rodríguez, and Juan Rojas for their invaluable contribution to the fieldwork for this project. Our gratitude also goes to the Smithsonian Tropical Research Institute and the research station at Coibita/Ranchería for support of the fieldwork.

