## Supplementary material for "White-faced capuchins (*Cebus capucinus imitator*) exhibit selectivity for stone tools": SI Appendix

**This PDF file includes:**

Tables S1 to S3

**Other supporting materials for this manuscript include the following:**

Movies S1 to S4

Tables

Table S1. Metric dimensions of locally available raw materials and hammerstone categories by island and tool function.

| **COIBA** | ***n*** | **Med. weight (g)** | **Med. length (mm)** | **Med. width (mm)** | **Med. thick (mm)** |
| --- | --- | --- | --- | --- | --- |
| Raw Material | 63 | 380 | 92 | 66 | 40 |
| Hammerstone (*Astrocaryum*) | 66 | 870 | 123 | 87 | 58 |
| Hammerstone (*Clypeolum*) | 51 | 180 | 69 | 50 | 34 |

| **JICARÓN** | ***n*** | **Med. weight (g)** | **Med. length (mm)** | **Med. width (mm)** | **Med. thick (mm)** |
| --- | --- | --- | --- | --- | --- |
| Raw Material | 202 | 155 | 88 | 63 | 37 |
| Hammerstone (*Terminalia*) | 261 | 675 | 121 | 91 | 53 |
| Hammerstone (*Coenobita*) | 170 | 400 | 107 | 78 | 46 |

Table S2. Parameter summaries of posterior mean, standard deviation (sd) and 89% highest posterior density interval (HPDI) of mean (α) and scale (φ) parameters from GLMs. Subscripts indicate raw material (rm) , *Terminalia catappa* (tc) , and *Coenobita compressus* (cc) and the dimensions of weight (w), thickness (th), length (l), and width (w) in the assemblages from Jicarón.

| parameter | mean | sd | 5.5% HPDI | 94.5% HPDI |
| --- | --- | --- | --- | --- |
| α_rm_wt | 5.97 | 0.08 | 5.85 | 6.10 |
| α_cc_wt | 6.27 | 0.06 | 6.18 | 6.37 |
| α_tc_wt | 6.60 | 0.04 | 6.54 | 6.67 |
| φ_rm_wt | 503.07 | 55.88 | 420.66 | 598.95 |
| φ_cc_wt | 321.49 | 36.95 | 267.22 | 381.93 |
| φ_tc_wt | 338.93 | 29.97 | 294.24 | 389.46 |
| α_rm_th | 3.72 | 0.03 | 3.68 | 3.77 |
| α_cc_th | 3.85 | 0.02 | 3.81 | 3.89 |
| α_tc_th | 3.98 | 0.02 | 3.96 | 4.01 |
| φ_rm_th | 7.64 | 0.78 | 6.47 | 8.95 |
| φ_cc_th | 4.57 | 0.50 | 3.85 | 5.41 |
| φ_tc_th | 4.65 | 0.41 | 4.04 | 5.34 |
| α_rm_l | 4.52 | 0.03 | 4.48 | 4.57 |
| α_cc_l | 4.70 | 0.03 | 4.65 | 4.74 |
| α_tc_l | 4.82 | 0.02 | 4.79 | 4.85 |
| φ_rm_l | 15.83 | 1.66 | 13.36 | 18.64 |
| φ_cc_l | 12.03 | 1.32 | 10.07 | 14.32 |
| φ_tc_l | 10.78 | 0.97 | 9.31 | 12.42 |
| α_rm_wd | 4.19 | 0.03 | 4.14 | 4.24 |
| α_cc_wd | 4.36 | 0.02 | 4.32 | 4.40 |
| α_tc_wd | 4.50 | 0.02 | 4.47 | 4.52 |
| φ_cc_wd | 8.22 | 0.90 | 6.90 | 9.77 |
| φ_tc_wd | 6.92 | 0.62 | 6.02 | 7.93 |
| φ_rm_wd | 12.16 | 1.24 | 10.32 | 14.33 |

**Table S3.** Parameter summaries of posterior mean, standard deviation (sd) and 89% highest posterior density interval (HPDI) of mean (α) and scale (φ) parameters from GLMs. Subscripts indicate raw material (rm) , *Astrocaryum standleyanum* (as) , and *Clypeolum latissimum* (fs) and the dimensions of weight (w), thickness (th), length (l), and width (w) in the assemblages from Coiba.

| **parameter** | **mean** | **sd** | **5.5% HPDI** | **94.5% HPDI** |
| --- | --- | --- | --- | --- |
| α_rm_wt | 6.31 | 0.1 | 6.15 | 6.48 |
| α_fs_wt | 5.42 | 0.11 | 5.24 | 5.6 |
| α_as_wt | 6.82 | 0.07 | 6.71 | 6.93 |
| φ_rm_wt | 400.43 | 71.83 | 299.02 | 525.03 |
| φ_fs_wt | 149.61 | 31.43 | 106.9 | 203.18 |
| φ_as_wt | 316.73 | 53.93 | 241 | 411.85 |
| α_rm_th | 3.8 | 0.05 | 3.72 | 3.88 |
| α_fs_th | 3.53 | 0.05 | 3.45 | 3.6 |
| α_as_th | 4.1 | 0.03 | 4.04 | 4.15 |
| φ_rm_th | 7.55 | 1.44 | 5.55 | 10.06 |
| φ_fs_th | 4 | 0.83 | 2.89 | 5.46 |
| φ_as_th | 4.53 | 0.83 | 3.38 | 5.97 |
| α_rm_l | 4.58 | 0.04 | 4.51 | 4.65 |
| α_fs_l | 4.24 | 0.05 | 4.16 | 4.31 |
| α_as_l | 4.8 | 0.03 | 4.76 | 4.84 |
| φ_rm_l | 10.75 | 2 | 8.03 | 14.36 |
| φ_fs_l | 7.31 | 1.56 | 5.23 | 9.99 |
| φ_as_l | 5.58 | 1.05 | 4.14 | 7.39 |
| α_rm_wd | 4.26 | 0.04 | 4.19 | 4.33 |
| α_fs_wd | 3.97 | 0.05 | 3.9 | 4.05 |
| α_as_wd | 4.46 | 0.03 | 4.41 | 4.5 |
| φ_fs_wd | 6.18 | 1.3 | 4.39 | 8.47 |
| φ_as_wd | 4.53 | 0.82 | 3.42 | 5.95 |
| φ_rm_wd | 8.72 | 1.6 | 6.44 | 11.5 |

Movie S1 (separate file). Video footage of hammerstone use on *Astrocaryum* by wild capuchins on Isla Coiba, Panama.

DOI: 10.5281/zenodo.17643935

Movie S2 (separate file). Video footage of hammerstone use on *Coenobita* hermit crabs by wild capuchins on Isla Jicarón, Panama.

DOI: 10.5281/zenodo.17643935

Movie S3 (separate file). Video footage of hammerstone use on *Clypeolium* freshwater Nerite snails by wild capuchins on Isla Coiba, Panama.

DOI: 10.5281/zenodo.17643935

Movie S4 (separate file). Video footage of hammerstone use on *Terminalia* nuts by wild capuchins on Isla Jicarón, Panama.

DOI: 10.5281/zenodo.17643935
